# Transformation of meiotic drive into hybrid sterility in Drosophila

**DOI:** 10.1101/2024.05.10.593569

**Authors:** Jackson Bladen, Hyuck-Jin Nam, Nitin Phadnis

**Affiliations:** School of Biological Sciences, University of Utah, Salt Lake City, UT 84112, USA

**Keywords:** meiotic drive, hybrid incompatibility, Sex-Ratio, *Drosophila*, speciation, genomic conflict

## Abstract

Hybrid male sterility is one of the fastest evolving intrinsic reproductive barriers between recently isolated populations. A leading explanation for the evolution of hybrid male sterility involves genomic conflicts with meiotic drivers in the male germline. There are, however, few examples directly linking meiotic drive to hybrid sterility. Here, we report that the *Sex-Ratio* chromosome of *Drosophila pseudoobscura*, which causes X-chromosome drive within the USA subspecies, causes near complete male sterility when moved into the genetic background of the Bogota subspecies. In addition, we show that this new form of sterility is genetically distinct from the sterility of F1 hybrid males in crosses between USA males and Bogota females. Our observations provide a tractable study system where non-cryptic drive within species is transformed into strong hybrid sterility between very young subspecies.

## Introduction

Hybrid male sterility is one of the fastest-evolving intrinsic reproductive barriers between geographically isolated populations (Wu 1992; Wu and Davis 1993; Coyne and Orr 1997). This rapid evolution of hybrid male sterility is thought to be driven by intra-genomic conflicts, particularly those involving meiotic drivers (Frank 1991; Hurst and Pomiankowski 1991; Henikoff *et al*. 2001; Tao and Hartl 2003). Meiotic drivers are selfish chromosomes that eliminate gametes that carry competing homologous chromosomes and thus over-transmit themselves above Mendelian expectations (Gershenson 1928; Sandler and Novitski 1957). These drivers can rapidly spread through populations due to their selfish transmission and, in turn, can be silenced by the rapid evolution of suppressors of drive (Hamilton 1967; Jaenike 2001; Hall 2004). When a driver is silenced, normal Mendelian segregation is restored and a history of meiotic drive becomes difficult to detect; such suppressed drive systems are therefore referred to as “cryptic drive systems” (Tao *et al*. 2001; Orr and Irving 2005; Presgraves and Meiklejohn 2021). The unmasking of such cryptic drive systems in hybrids is central to our understanding of the observed rapid evolution of male sterility between recently diverged populations (Frank 1991; Hurst and Pomiankowski 1991).

There are at least two ways that unmasking of cryptic drive systems may lead to hybrid male sterility. Under the first version, which we refer to as the “crossfire” model, when two populations of a species are geographically isolated, each population may go through its private history of accumulating cryptic drivers. When these two populations hybridize, multiple cryptic drivers accumulated in each parental lineage may become simultaneously unmasked in a naïve hybrid genetic background. A cross-firing of multiple drive systems in hybrids could eliminate all gametes and cause sterility. Under the second version, which we refer to as the “misfire” model, a single cryptic drive system in one population may be sufficient to explain the evolution of hybrid male sterility. When a cryptic drive system is unmasked in hybrids, it may misfire and destroy all gametes rather than destroying only targeted gametes, causing hybrid male sterility. Instead of simply re-manifesting as drive upon unmasking, this misfiring of drive necessitates a genetic background-dependent transformation of drive within a population into sterility between populations.

The best example of the “crossfire” model comes from within-species sterility of hybrids between isolates of the fission yeast *Schizosaccharomyces pombe*, which are sterile due to the unmasking of multiple *wtf* meiotic drive genes (Zanders *et al*. 2014; Hu *et al*. 2017; Nuckolls *et al*. 2017; Bravo Núñez *et al*. 2020). There are, however, no examples where inter-species hybrids are known to be sterile due to the unmasking of multiple meiotic drivers. The best example of the “misfire” model comes from inter-species hybrids between the Bogota and USA subspecies of *Drosophila pseudoobscura* (Orr and Irving 2005; Phadnis and Orr 2009). F1 hybrid males from crosses between Bogota mothers and USA fathers are sterile (Prakash 1972) but recover weak fertility when aged and show *X*-chromosome segregation distortion (Orr and Irving 2005). Here, hybrid male sterility and *X*-chromosome meiotic drive have a shared genetic basis (Orr and Irving 2001, 2005; Phadnis 2011) and at least one gene, *Overdrive*, is required for both phenotypes (Phadnis and Orr 2009). Genetic analyses are consistent with the idea that unmasking a single cryptic Bogota *X*-chromosome driver causes hybrid male sterility; there is no evidence for multiple meiotic drivers in these hybrids.

Another line of evidence supporting the “misfire” model involves asking whether known meiotic drivers within populations can cause sterility when crossed into a naïve genetic background. The best examples of this approach involve studies of *X*-chromosome meiotic drivers (*Sex-Ratio* chromosomes) in *D. subobscura* (Hauschteck-Jungen 1990; Verspoor *et al*. 2018) and *D. simulans* (Merçot *et al*. 1995; Cazemajor *et al*. 1997). In these species, the unmasking of locally suppressed *Sex-Ratio* chromosomes leads to a reduction in male fertility when crossed to strains that do not carry suppressors-of-drive (Hauschteck-Jungen 1990; Merçot *et al*. 1995; Cazemajor *et al*. 1997; Verspoor *et al*. 2018). However, the reduction in male fertility in the above cases is modest and does not appear substantially different from that expected from the reduction in the number of sperm due to drive alone (Montchamp-Moreau and Joly 1997; Verspoor *et al*. 2018).

Here, we report that the *Sex-Ratio* chromosome of the USA subspecies of *D. pseudoobscura* causes near complete hybrid male sterility when moved into an otherwise Bogota genetic background. This new form of hybrid male sterility is genetically distinct from the sterility of hybrid F1 males in crosses between Bogota mothers and USA fathers. Our results show that even a single unsuppressed meiotic driver is sufficient to cause male sterility when moved into a naïve genetic background. Although the role of meiotic drive-related genomic conflicts in the evolution of hybrid sterility is a long-debated topic in biology, the lack of suitable biological systems to study these phenomena remains a key limiting factor. Our discovery provides a strong study system to understand how a single meiotic driver within species can transform into male sterility between species.

## Results

The *D. pseudoobscura Sex-Ratio* (*SR*) chromosome is an *X*-chromosome meiotic driver segregating in the USA subspecies of *D. pseudoobscura* (Sturtevant and Dobzhansky 1936). The USA *SR* chromosome is an unsuppressed drive system; no suppressors against this *SR* chromosome have been found despite repeated surveys of natural populations since its original discovery more than 85 years ago (Sturtevant and Dobzhansky 1936; Policansky and Dempsey 1978; Beckenbach *et al*. 1982; Price *et al*. 2014). This *SR* chromosome carries three non-overlapping chromosomal inversions on the right arm of the metacentric *X*-chromosome (*XR*) (Sturtevant and Dobzhansky 1936; Fuller *et al*. 2020). The genetic basis of meiotic drive in this *SR* chromosome is thought to be complex (Wu and Beckenbach 1983) and all genes necessary for drive are located on the inversion-bearing *XR*.

USA and Bogota are very young subspecies in the earliest stages of divergence and are estimated to have been geographically separated for only ∼155-230K years (Schaeffer and Miller 1991; Machado *et al*. 2002). The USA *SR* chromosome, in contrast, is ancient and is estimated to be ∼1 million years old (Babcock and Anderson 1996; Fuller *et al*. 2020). Although the Bogota subspecies harbors some of the chromosomal inversion polymorphisms found in the USA subspecies, the *SR* chromosome is not found in Bogota (Dobzhansky *et al*. 1963). The *SR* chromosome has been present in the ancestor of Bogota and USA for most of its 1 million years existence, except for the last 150-230K years, where it has only existed in the USA population. If Bogota has evolved suppressors against the *SR* chromosome, this could explain the absence of *SR* in this subspecies.

To test whether Bogota harbors dominant suppressors against *SR*, we generated F1 hybrid males that carry the USA *SR* chromosome. Hybrid F1 males between USA mothers and Bogota fathers are fertile and show normal segregation patterns (Prakash 1972; Orr and Irving 2005). We crossed USA females carrying the *SR X*-chromosome with Bogota males from several strains. The resulting F1 hybrid males, which carry the USA *SR* chromosome in a hybrid autosomal background, are fertile and produce nearly 100% female progeny. The average proportion of female offspring produced by F1 hybrid males carrying the USA *SR* chromosome, across six Bogota strains, was 97% females. The average proportion of female offspring produced by F1 hybrid males carrying the USA *ST* chromosome, across six Bogota strains, was 46% females. These results confirm that Bogota does not carry dominant suppressors of the USA *Sex-Ratio* chromosome, consistent with previous findings (Orr and Irving 2005).

Next, to test whether Bogota harbors recessive suppressors against the *SR* chromosome, we introgressed the USA *SR* chromosome into an otherwise homozygous Bogota background. We performed this introgression through repeated backcrossing to Bogota while selecting for the *SR* chromosome in each generation. Because there are no visible genetic markers to assist with this introgression, we first developed PCR-based markers that detect all three *SR*-associated inversions. Because recombination is suppressed along the whole *SR* chromosome (Fuller *et al*. 2020), tracking our introgression with a single molecular marker is sufficient to move the entire *SR* chromosome arm into Bogota. We used a PCR-based marker on the *Basal* inversion to select for *SR* chromosome-bearing females in each generation for 10 generations of repeated backcrosses with Bogota males (Figure 1). The *USA SR* chromosome arm is thus maintained in this crossing scheme while the remaining genetic background becomes increasingly Bogota with each backcross generation.

**Figure 1.**
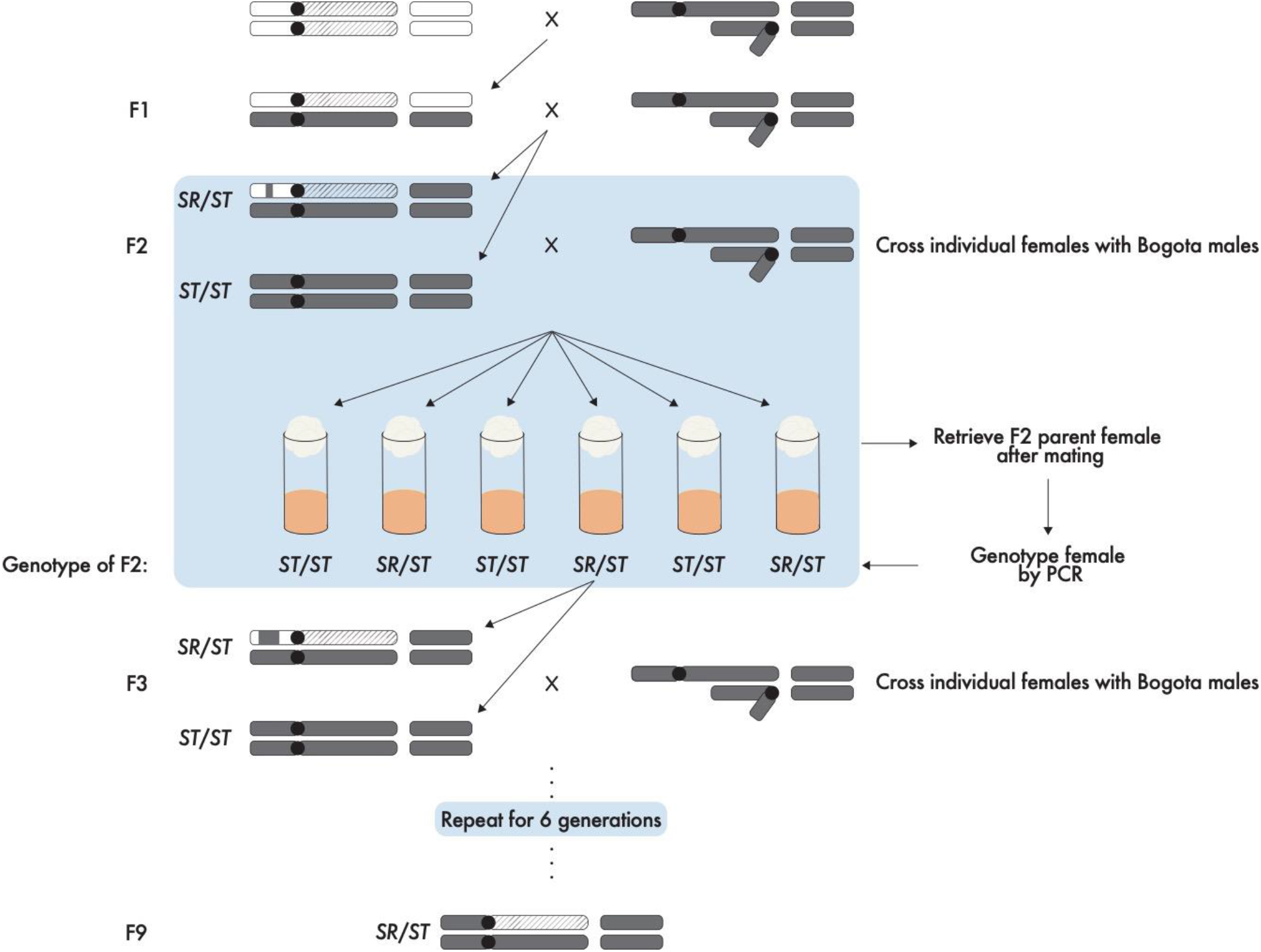
Crossing scheme for the introgression of the USA *SR* chromosome into a Bogota genetic background. Introgression scheme to generate introgression males carrying the *SR* chromosome in a largely Bogota genetic background. Starting with the F2 generation, females were backcrossed individually to Bogota males. After mating, females were collected from crosses and genotyped using PCR to identify whether they carried the *SR* polymorphism or the Bogota *X*-chromosome. If the female carried the *SR* chromosome, her female offspring of unknown genotype were backcrossed individually to Bogota males. This process was independently repeated for nine generations with three Bogota strains. The *SR* chromosome remains unaltered through the introgression procedure because it suppresses recombination along the entire *XR* chromosome arm. USA chromosomes are shown with empty ovals, USA *SR* chromosomes with diagonally lined ovals, and Bogota chromosomes with dark gray ovals. Only the sex chromosomes and second autosomes are shown for clarity.

At the end of the introgression procedure, females in this cross are heterozygous at the right arm of the *X*-chromosome, carrying one copy of USA *SR* and one copy of Bogota *XR* in an otherwise Bogota genetic background (Figure 2A). When these females are crossed with Bogota males, two types of sons are produced. The first type of sons are pure Bogota males. As expected, pure Bogota sons produced from this cross are fertile and show normal Mendelian segregation (Figure 2B). The second type of sons are identical to pure Bogota males, except they carry USA *SR* on *XR* (Figure 2A). Surprisingly, we found that males carrying USA *SR* in a Bogota background are sterile (Figure 2B). Normally, males carrying a USA *Standard* (*ST*) *XR* in a Bogota background are fertile (Phadnis and Orr 2009). Here, we tested 83 *SR* introgression males across three different Bogota genetic backgrounds. Almost none of them produced any progeny; only three exceptional males produced fewer than four progeny. Simply substituting *ST* for *SR* – even though they both represent USA material on *XR* – results in male sterility in a Bogota genetic background. The phenotype of the *SR* chromosome thus appears to transform from meiotic drive in a USA background into male sterility in a Bogota background.

**Figure 2.**
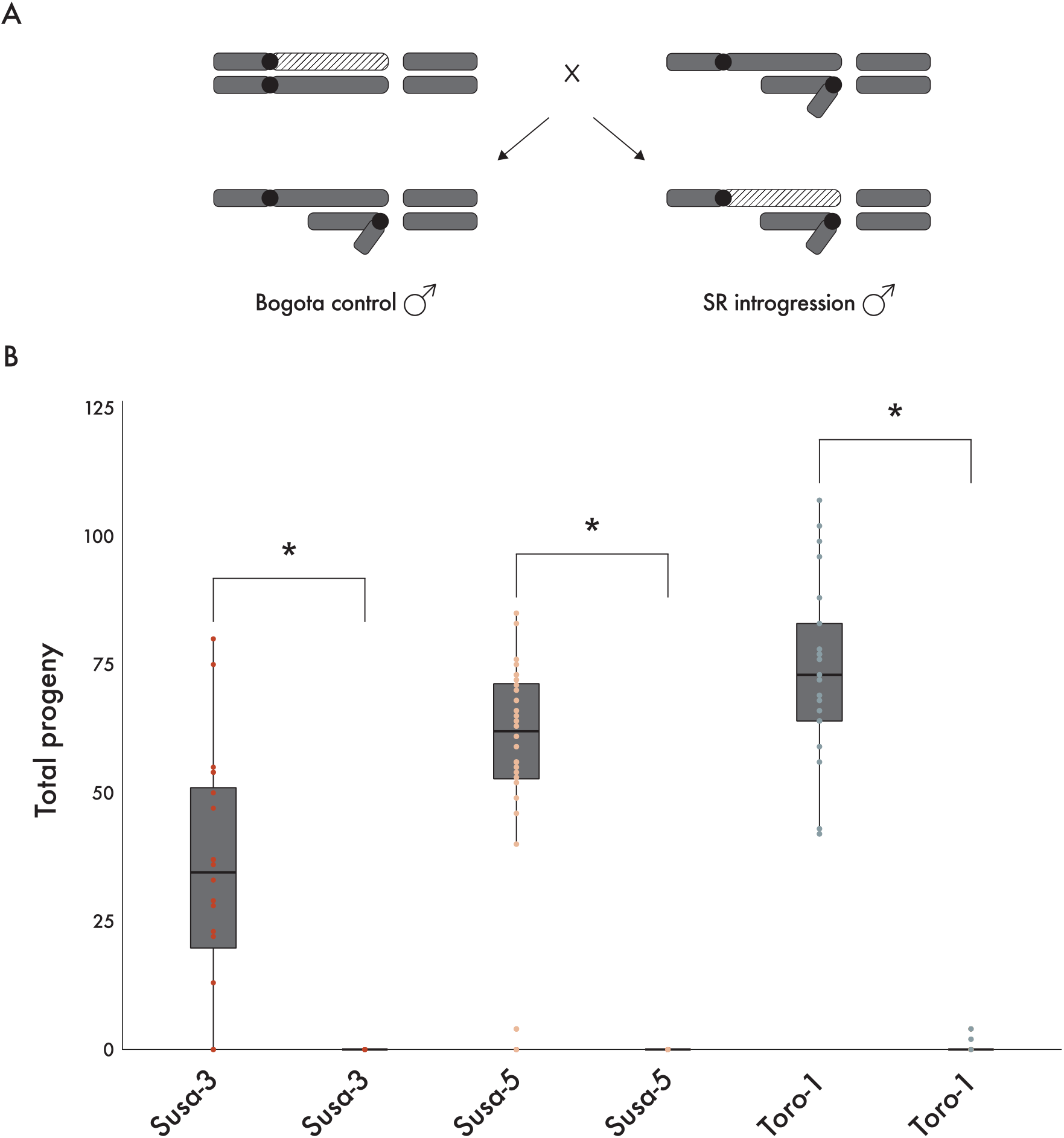
The USA *SR* chromosome causes near-complete sterility in a Bogota genetic background. A. After nine generations of backcrossing with Bogota males, *SR* introgression females were crossed with Bogota males. These crosses produce two types of sons. The first type of son is a pure species Bogota male (Bogota control male); the second type of son is identical to the first except that it carries the USA *SR* on *XR (SR* introgression male). These males, of unknown genotype, were crossed individually with Bogota females from the Bogota strain used during the introgression procedure. After mating, males were collected from crosses and genotyped using PCR to identify whether they carried the *SR* polymorphism or the Bogota *X*-chromosome. Bogota chromosomes are shown with dark gray ovals and the USA *SR* chromosome arm is shown with diagonally striped ovals. B. Total progeny counts of Bogota control males and *SR* introgression males from three Bogota strains. Data points corresponding to Bogota control males and *SR* introgression males from the same Bogota strain share a color. Progeny counts from Bogota control males are shown with dark gray boxes (Susa-3, *n* = 20; Susa-5, *n* = 28; Toro-1, *n* = 21). Progeny counts from *SR* introgression males are shown by the absence of boxes (Susa-3, *n* = 19; Susa-5, *n* = 38; Toro-1, *n* = 26). Floating bars with an asterisk above indicate treatments that were significantly different from each other (Pairwise Wilcoxon Rank Sum test, *p* < 1.80e-6 for all comparisons).

The *SR* chromosome is recombinationally inert when heterozygous with the *ST* chromosome even in large colinear regions spanning several megabases (Fuller *et al*. 2020). Although the reasons for this near-complete suppression of recombination outside of inversions remain mysterious, recombination between inversions can occur at a low, non-zero rate (Wallace 1948; Fuller *et al*. 2020). If the observed male sterility is an artifact of some incidental alteration of the *SR* chromosome during the introgression procedure, then it may no longer be capable of causing meiotic drive even when re-introduced into an F1 hybrid background. If the introgressed *SR* chromosome remained intact, however, then the male sterility caused by *SR* is predicted to revert to *X*-chromosome meiotic drive when re-introduced into an F1 hybrid background. To test this idea, we crossed the heterozygous introgression females that carry one copy of Bogota *ST* and one copy of the USA *SR* with USA males (Figure 3). These crosses produce two types of sons. The first type of sons are reconstituted F1 hybrid males from crosses between Bogota mothers and USA fathers (Figure 3A). As expected, these sons produced almost no progeny (Figure 3B). The second type of sons are identical to the first type, except they carry USA *SR* on *XR* (Figure 3A). These sons were fertile and produced nearly 100% daughters (Figure 3B). This reversion of phenotype from hybrid male sterility back to *X*-chromosome meiotic drive shows that the introgressed *SR* chromosome remained intact and capable of causing drive even after 10 generations of backcrosses. We conclude that meiotic drive caused by the USA *SR* chromosome within species transforms into male sterility between species.

**Figure 3.**
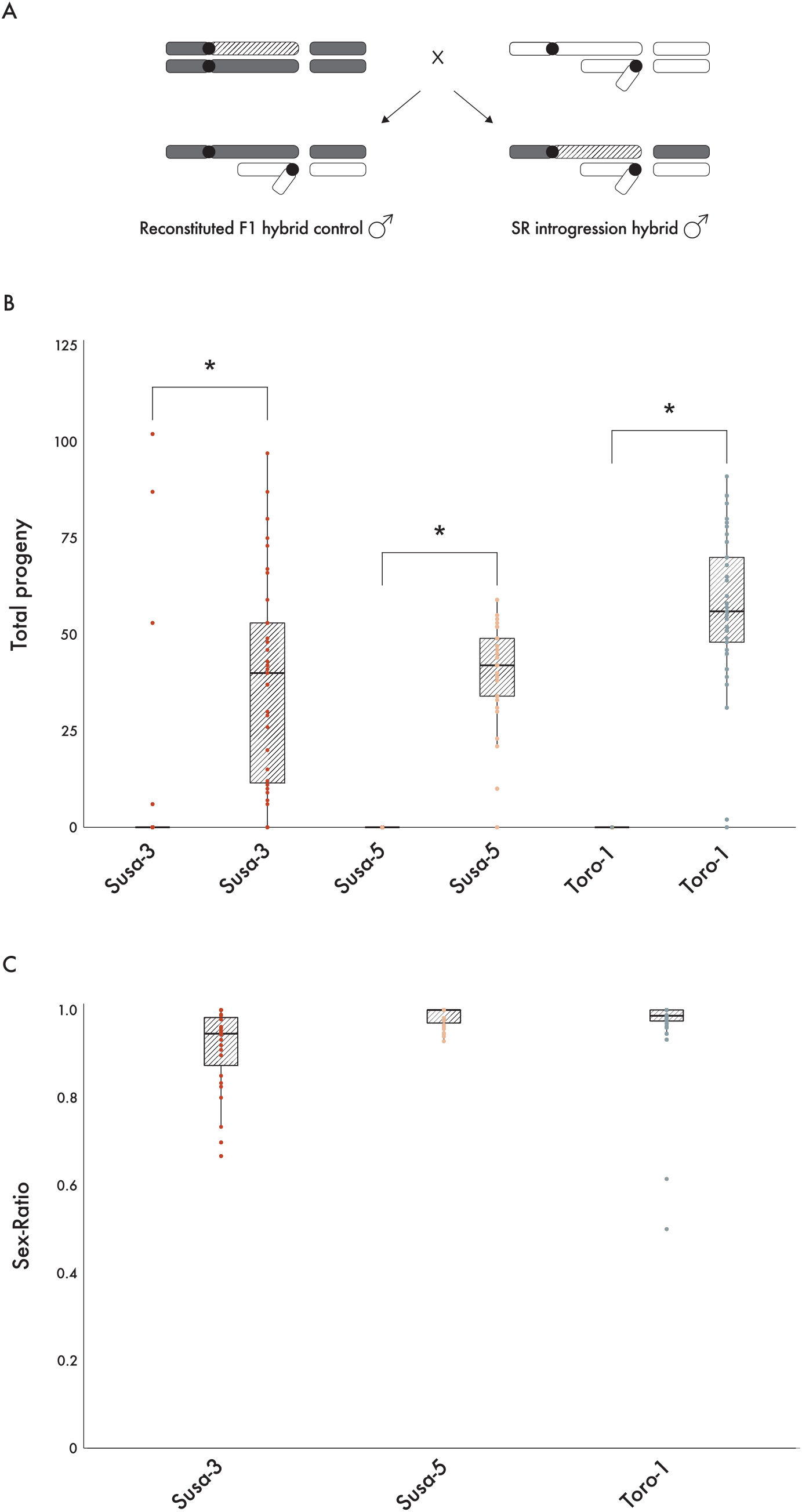
The USA *SR* chromosome remains capable of distortion after 10 generations of backcrossing. A. After nine generations of backcrossing with Bogota males, *SR* introgression females were crossed with USA males. These crosses produce two types of sons. The first type of son is a reconstituted F1 hybrid control male, nearly identical to those produced from crosses between Bogota females and USA males. The second type of son is identical to the first type, except it carries USA *SR* on *XR* (*SR* introgression hybrid male). These hybrid males, of unknown genotype, were crossed individually with Bogota females from the respective Bogota strain used during the introgression procedure. After mating, hybrid males were collected from crosses and genotyped using PCR to identify whether they carried the *SR* polymorphism or the Bogota *X*-chromosome. USA chromosomes are shown with empty ovals, USA *SR* chromosomes with diagonally lined ovals, and Bogota chromosomes with dark gray ovals. B. Total progeny counts of reconstituted F1 hybrid control males and *SR* introgression hybrid males. Data points corresponding to reconstituted F1 hybrid control males and *SR* introgression hybrid males from the same Bogota strain share a color. Progeny counts from reconstituted F1 hybrid control males are shown by the absence of boxes (Susa-3, *n* = 38; Susa-5, *n* = 35; Toro-1, *n* = 46). Progeny counts from *SR* introgression hybrid males are shown with diagonally lined boxes (Susa-3, *n* = 35; Susa-5, *n* = 29; Toro-1, *n* = 41). Floating bars with an asterisk above indicate treatments that were significantly different from each other (Pairwise Wilcoxon Rank Sum test, p < 3.08e-9 for all comparisons). C. Progeny sex-ratios of *SR* introgression hybrid males from each Bogota strain. Sex-ratio is calculated as the proportion of female progeny produced by each male. Sex-ratio values were not calculated if a male produced fewer than ten offspring.

## Discussion

The Bogota and USA subspecies of *D. pseudoobscura* are considered paradigmatic of the earliest stages of speciation (Lewontin 1974). Previous work has largely focused on understanding the genetic basis of male sterility and meiotic drive in F1 hybrid males from crosses between Bogota mothers and USA fathers (Prakash 1972; Dobzhansky 1974; Orr 1989a; b; Orr and Irving 2001, 2005; Phadnis and Orr 2009; Phadnis 2011). This F1 hybrid male sterility between Bogota mothers and USA fathers involves a complex interaction between factors on the left and right arms of the Bogota *X*-chromosome and *dominant* USA autosomal factors (Prakash 1972; Orr and Irving 2001; Phadnis 2011). In contrast, the sterility of the USA *SR* chromosome in an otherwise Bogota genetic background involves an interaction between factors on the USA *Sex-Ratio XR* and *recessive* Bogota autosomal factors. This new form of *SR*-induced sterility that we describe here thus appears genetically unrelated to F1 hybrid male sterility between Bogota mothers and USA fathers.

Previous explanations for the evolution of hybrid male sterility have focused on the unmasking of cryptic drive systems. The *D. pseudoobscura SR* chromosome is not a cryptic drive system; no suppressors of *SR* have been identified. Yet, a cryptic *suppressor* may potentially explain our observations. The USA *SR*-induced sterility described here must result from evolutionary changes accumulated after the USA-Bogota split. Consider a scenario where the *SR* chromosome – after this split – has gone through one or more bouts of suppression followed by escape from suppression in the USA population. The current unsuppressed state of the *SR* chromosome may thus represent a transient state in an evolutionary arms race where it currently holds the upper hand. When re-introduced to a naïve Bogota genetic background, an imbalance between drive and the missing suppressors may lead to the indiscriminate destruction of all gametes. A cryptic suppression system may thus better explain hybrid male sterility caused by the *SR* chromosome in a Bogota background.

The transformation of meiotic drive into sterility observed here necessitates a mechanism where the driving *SR*-chromosome – in addition to destroying *Y*-chromosomes – gains a new property of also destroying itself. Drivers that destroy themselves, known as ‘*suicide chromosomes*’, have been isolated in the *Segregation Distorter* system in *D. melanogaster* (Sandler and Hiraizumi 1960; Hartl 1974; Larracuente and Presgraves 2012). These suicide chromosomes, however, are artificially generated recombinant chromosomes where the target of drive is moved onto the driving chromosome. This scenario does not explain our observed transformation of drive into sterility because recombination is absent between sex chromosomes. Other natural cases of genetic background-dependent change in the *direction* of *Sex-Ratio* drive have been described in *D. affinis*. In *D. affinis*, nullo-*Y* males are fertile (Voelker and Kojima 1971). When the *D. affinis SR* chromosome is present in a nullo-*Y* male, it destroys itself and produces a male-biased progeny sex ratio. This phenomenon is known as *Male Sex Ratio* (*Msr*) (Novitski 1947; Voelker 1972). Although the mechanisms of this ‘drive rebound’ remain unknown, the *Sex-Ratio* chromosomes in an *Msr* genetic background can destroy gametes that carry themselves (Ma *et al*. 2022). Cases of drive rebound, however, remain fertile despite a reversal of the direction of drive and thus cannot fully explain our observed *SR*-induced sterility.

A mechanistic framework analogous to a toxin-antidote system may better explain our observations. Consider an *X*-chromosome driver that produces a toxin to which both *X*- and *Y*-bearing gametes are susceptible to differing degrees. If *Y*-bearing gametes are more susceptible to this toxin than *X*-bearing gametes, this can generate *X*-chromosome drive. The evolution of suppressors of drive may act as an antidote to this toxin. An evolutionary arms race between drive and suppression may involve a gradual increase in dosage of the toxin balanced by an increase in dosage of the antidote through stepwise co-evolution. In a hybrid background where the dosage of the toxin and antidote is imbalanced, the excessive toxin may destroy both *X*- and *Y*-bearing gametes and cause male sterility.

Our study shows that the theoretical minimum number of drivers – one – is sufficient to cause hybrid male sterility between species. Whether the driver or the suppressor is cryptic within a population appears to not matter for the manifestation of hybrid male sterility between populations. The USA *SR*-induced sterility in a Bogota genetic background provides a strong case study for the misfire model of the rapid evolution of hybrid male sterility in a tractable system where drive within species is transformed into hybrid sterility between species. Our ongoing studies to identify the causal genes and to understand the molecular and developmental mechanisms of USA *SR*-induced sterility in a Bogota background may shed light on how drive systems that have evolved to selectively destroy targeted gametes may inadvertently destroy all gametes in hybrids, thus contributing to the origins of new species.

## Material and methods

### Fly strains and culture conditions

The *SR* chromosome was identified from natural population collections in Zion National Park, Utah, in September 2013. The base *SR* stock was created by re-isolation of the *SR* chromosome and seven generations of backcrossing to an inbred stock with the *Standard* arrangement *X*-chromosome carrying visible mutations *sepia* and *short* on the right arm of the *X*-chromosome. The resulting *SR* chromosome stock segregates for *SR* and *Standard X*-chromosomes, the latter being identified by visible mutations. For a comprehensive list of all other fly strains used in this study, see Table 1. All experimental crosses were performed on food containing standard cornmeal molasses *Drosophila* media seeded with live yeast. After the adults were removed from the crosses, the food was hydrated as needed with 0.5% v/v propionic acid. Unless stated otherwise, all crosses were maintained at room temperature on a 14:10 hour light-dark cycle.

**Table 1.** List of fly strains used in this study.

| Name | Species | Origin |
| --- | --- | --- |
| Sex-Ratio | USA | Phadnis Lab |
| *w*<sup>152</sup> | USA | Drosophila Species Stock Center |
| *se*<sup>1</sup>, *sh*<sup>1</sup> | USA | Drosophila Species Stock Center |
| *Susa-2* | Bogota | Drosophila Species Stock Center |
| *Susa-3* | Bogota | Drosophila Species Stock Center |
| *Susa-5* | Bogota | Drosophila Species Stock Center |
| *Toro-1* | Bogota | Drosophila Species Stock Center |
| *Toro-4* | Bogota | Drosophila Species Stock Center |
| *Potosi-1* | Bogota | Drosophila Species Stock Center |

### Test for dominant suppressors

To test whether the Bogota population carries dominant suppressors for the *SR* chromosome, we crossed USA females heterozygous for the *SR* polymorphism with Bogota males from 5 different inbred Bogota strains – *Susa-2, Susa-3, Susa-5, Toro-4*, and *Potosi-1*. Each cross consisted of 10 unmated USA *SR* females and 10 Bogota males. These crosses produced F1 hybrid males hemizygous for the *SR* polymorphism in a hybrid autosomal background. The F1 hybrid males from each of the five treatments were individually mated to virgin females from USA *white*. After all flies had eclosed, we scored the sex of the F2 progeny. The sex-ratio phenotype was measured as the count of female offspring divided by the total number of offspring. For statistical analyses, crosses producing greater than 10 total offspring were used for calculating sex-ratio.

### Introgression procedure

We first crossed USA females homozygous for the *SR* polymorphism with Bogota males. All F1 hybrid females were heterozygous for the *SR* polymorphism. We backcrossed these F1 hybrid females with Bogota males to generate F2 females, which were either heterozygous for the *SR* polymorphism or homozygous for the Bogota *X*-chromosome. We randomly selected 10-15 of these virgin females and crossed each female independently with three Bogota males. After 5-7 days of mating, we disposed of the males and retrieved the female from each cross. We performed PCR on the genomic DNA from each female to identify which F2 females carried the *SR* polymorphism. When PCR confirmed the F2 mother carried the *SR* polymorphism, we selected 10-15 of her unmated female progeny (F3) and crossed each female independently with three Bogota males. We repeated this procedure for a total of ten generations. The introgression procedure was performed using three strains caught within Bogota, Columbia - *Susa-3, Susa-5*, and *Toro-1*. To track the *SR* chromosome through the introgression procedure, we designed PCR primers that amplify DNA within the *Basal* inversion of *SR* – F primer: TCTTATCAAAGGGATTGACC, R primer: CCTATGTGGACATCATCTTT. Importantly, these primers do not amplify DNA from any of the USA or Bogota strains used in this study.

After ten generations of backcrossing with Bogota males, we randomly selected introgression males from each treatment and crossed each male individually with virgin Bogota females from the respective Bogota strain. The introgression males either carried the USA *XR Sex-Ratio* chromosome in a largely Bogota genetic background or the Bogota *X*-chromosome in a largely Bogota genetic background. After 7 days of mating, we disposed of the females and retrieved the male from each cross. We used our PCR-based molecular marker to identify whether males carried the *SR* polymorphism or the Bogota *X*-chromosome. We scored the sex of the F2 progeny after all flies had eclosed.

### Validating introgression procedure

To test whether the *SR* chromosome was altered during the introgression procedure, we crossed F9 introgression females, who carry one copy of the USA *SR* chromosome in a largely Bogota genetic background, with USA *white* males. We randomly selected F1 males from each introgression strain and crossed them individually with virgin Bogota females from the respective Bogota strain used during the introgression procedure. After 7 days of mating, we disposed of the females and retrieved the male from each cross. We used our PCR-based molecular marker to identify whether males carried the *SR* polymorphism or the Bogota *X*-chromosome. We scored the sex of the F2 progeny after all flies had eclosed.

## Acknowledgements

We are grateful to Jackson Ridges and Rob Unckless for helpful discussions and feedback in improving this manuscript. This work was supported by the National Institute of Health grants 5T32GM141848-3 to JB and R01GM141422 to NP. Strains are available upon request. The authors affirm that all data necessary for confirming the conclusions of the article are present within the article, figures, and tables.

